# Detecting Foldback Artifacts in Long-reads

**DOI:** 10.1101/2025.07.15.664946

**Authors:** Jakob M. Heinz, Matthew Meyerson, Heng Li

**Affiliations:** Department of Biomedical Informatics, Harvard Medical School, Boston, MA, United States; Department of Data Science, Dana-Farber Cancer Institute, Boston, MA, United States; Cancer Program, Broad Institute of MIT and Harvard, Cambridge, MA, USA; Department of Medical Oncology, Dana-Farber Cancer Institute, Boston, MA, USA; Department of Genetics, Harvard Medical School, Boston, MA, USA

**Keywords:** Long-read sequencing, Nanopore, RNA-Sequencing, Technical artifacts, Quality control

## Abstract

Long-read sequencing data is useful for detecting large and complex structural variations; however, technical artifacts can lead to false structural variant calls. In our analyses, we became aware of a foldback artifact in long-read data. Therefore, we developed the open-source Breakinator tool to flag putative foldback artifact reads, as well as previously known chimeric artifacts. Through an alignment-based approach, Breakinator can detect artifacts missed by existing quality control tools. We profiled the occurrences of foldbacks and chimeric reads in both Oxford Nanopore and PacBio sequences across a range of specimens, library types, sequencing chemistries, sequencing machines, and base-calling software.

## Background

Advances in long-read sequencing technologies, such as those by Pacific Biosciences (PacBio) and Oxford Nanopore Technologies (ONT), have enabled the reliable resolution of complex and repetitive regions of the genome, which cannot be achieved by short-read sequencing. The ability of long reads to span these complicated regions is useful for detecting large, complex structural variations (SV) in the genome [1, 2]. Typically, for long-read SV calling, reads are aligned to a reference genome to identify those with a supplementary alignment, a so-called split read, where a portion of the read maps to one genomic region, and the remainder maps to another distinct region, causing the aligner to split the read into separate alignment blocks [3–5]. However, systemic technical artifacts in the reads can lead to false SV calls. Chimeric reads, technical artifacts that concatenate two distinct sequences, have been a pervasive issue throughout the history of long-read sequencing [6–9]. Technical artifacts can arise through issues in library preparation or computational errors in distinguishing independent molecules [10, 11]

Here, we have become aware of another type of technical artifact, a foldback (also known as a duplicated inversion) [12], where half the read maps to one strand and the other half maps to the same location, but on the opposite strand, appearing as a near-perfect concatemerization of the forward and reverse strands of a read. Interestingly, previous ONT sequencing protocols, such as the 2D and 1D² protocols, intentionally generated foldback-like structures to enable the tandem sequencing of a template molecule and its complement to improve base-calling accuracy [13–15]. A previous study by Soneson *et al.* had noted an elevated rate of palindromic reads in ONT direct-cDNA data with supplementary alignments overlapping their primary alignment on the opposite strand. They hypothesized that these may represent un-split reads by the base caller [11]. These foldback artifacts were also noted in the recent somatic SV caller SAVANA by Elrick *et al.*, which includes a preprocessing step to remove putative foldback artifacts from tumors and matched germline controls before collecting SV supporting reads. The authors observed that foldback artifacts typically account for 1–4% of genomic (g)DNA reads that pass their filter across different tumor regions and patients, but could reach as high as 20–30% in some cases [16]. While foldback artifacts have been observed previously, to our knowledge, no studies have systematically characterized foldback artifacts across library types, sequencing chemistries, and sequencing machines using diverse samples. Therefore, we applied the open-source Breakinator tool we developed to profile foldback and the already well-known chimeric read artifacts in publicly available genomic and transcriptomic datasets, including recent sequencing technologies such as the ONT’s latest R10.4 nanopores and PacBio HiFi.

## Methods

All data used in this study were downloaded from publicly available sources, with the links available in Additional file 1. The K562 samples were generated by the SGN-Ex consortium [17]. HCC1395 samples were generated by Cotto *et al.* [18]. The mouse tissue samples were generated by Sessegolo *et al.* [19]. The HG002 gDNA samples were downloaded from ONT’s 2025 Genome in a Bottle Data (GIAB) Release [20]. The HG002 direct-RNA002, PacBio Iso-seq, and Mas-seq were made available by GIAB. The HG002 dRNA sample SRR30901279 and the HG002 cDNA data were generated by Zheng *et al.*[21] The HG002 PacBio HiFi gDNA is a public dataset made available by PacBio [22]. The PacBio scAAV sample was generated by Talevich *et al*. [23]

The samples were aligned to their respective reference genomes using Minimap2 [3]. K562 and HCC1395 samples were aligned to T2T-CHM13v2.0 [24, 25]. The mouse samples were aligned to GRCm39 [26]. HG002 samples were aligned to the HG002v1.0.1 diploid genome [27]. To align dRNA ONT, the following parameters were used: “-cx splice -uf -k14 --secondary=no”. For ONT direct-cDNA and cDNA alignments, “-cx splice --secondary=no” parameters were used. ONT gDNA data were aligned with “-cx map-ont --secondary=no” parameters. For PacBio Iso-Seq and Mas-Seq data, we used “-cx splice:hq -uf --secondary=no” parameters. Finally, PacBio HiFi data were aligned with “-cx map-pb -- secondary=no.” Aligning to a diploid genome decreases the mapping quality of many reads, as there will likely be at least two locations where they map well. To address this issue, HG002 sample reads were aligned to the HG002v1.0.1 diploid genome by aligning the reads to each haplotype separately using Minimap2 with the flags “--secondary=no” and “--paf-no-hit” specified. We wrote a preprocessing script for the Breakinator to compare the alignment of every read to the maternal and paternal haplotype assemblies, selecting the alignment with the highest alignment score as reported by the AS::i tag. If an alignment to one haplotype covered more than 95% of the read, it was chosen over a chimeric alignment to the other haplotype. If the alignment scores were equal, then the maternal haplotype alignment was chosen.

The PAF files were generated using either “paftools.js sam2paf -p” on a SAM file or Minimap2 with the “--secondary=no” flag. We designed the Breakinator to parse input SAM/BAM/CRAM format files or PAF format files to detect reads with supplementary alignments. Secondary alignments were ignored. Alignments with mapping quality less than 10 or lengths less than 200 bps were filtered out. The primary and all supplementary alignments for a given read were identified and sorted by the read start location of the alignment. If the first read segment mapped to the forward strand, then the end of the alignment of the first segment and the start of the second segment were identified as the breakpoint location. If the first segment mapped to the reverse strand, the reciprocal start and end locations were taken for the breakpoint. As shown in Fig. 1a, if the breakpoint was to the opposite strand within 200 bps of the end of the alignment of the first segment and the breakpoint occurs within a 10% margin on either side of the middle of the read, it was classified as a foldback. The default threshold of 200bps for foldback events was chosen to allow for potential ambiguity in the alignment borders by minimap2. If the breakpoint was to another chromosome (interchromosomal) or more than 1 Mb away (intrachromosomal), it was classified as a chimeric read. The 1 Mb default threshold for intrachromosomal chimeras was chosen because only two introns, out of 177,839, in MANE-selected protein-coding genes were longer than 1 Mb [28]. If the breakpoint was not chimeric or a foldback, then it was considered a true breakpoint. The identified breakpoints and their classification were generated for every sample. While uncommon, it is possible to have multiple breakpoints on the same read. To classify reads at the read level, we used the following approach: if a read contained any foldback or chimeric breakpoint, regardless of the number of true breakpoints, it was classified as an artifact. If there were multiple foldback or chimeric breakpoints on the read, the read was classified according to which breakpoint occurred most frequently. If they were equal, then the read was classified as chimeric. All summary statistics are generated using the read level classifications.

To compare how existing tools detect the reads we had identified as artifactual, we filtered all reads with fastplong, porechop, and restrander, respectively [29–31]. Fastplong was run with the following command: “fastplong -i sample.fastq -o filter_pass.fastq --failed_out failed_reads.fastq.” Restrander was run only on our cDNA samples, as it didn’t support any other data types, with the command “restrander sample.fastq filter_pass.fastq config.json.” Restrander required a configuration file of adaptor sequences, so for samples that did not have their library version supported by Restrander, we ran them with the closest version of the library supported by Restrander. For example, DCS108 samples were run with the SQK-DCS109 config file. Porechop was also only run on the transcriptomic data using the command: “porechop -i sample.fastq --discard_middle -o filter_pass.fastq.” For each tool, the read IDs that passed their respective filter were used to subset the reads that we found to pass our default filter, which requires a minimum of 200 bps of alignment and a mapping quality of at least 10. The reads classified as foldback, chimeric, or any breakpoint were similarly subset to include only the reads that passed the respective filter to calculate the proportions reported in Additional file 2. For the comparison, we ran Breakinator with the “–no-sym” parameter to capture all putative foldback events. PyChopper [32] was only run on the three HCC1395 samples, as we found it to be too slow to run on all samples. The length of the unaligned sequence between aligning segments of the breakpoint was identified by running Breakinator with the “--rcoord” flag, which outputs the read coordinates of the breakpoint along with the reference coordinates of the breakpoint. The reads were subset to foldback, chimeric, and true breakpoints, and the proportions of reads with at least 11 bps between the read coordinates of the breakpoint were calculated for each subset in Additional file 3. In the K562 samples, the coordinates of the unmapped sequences between aligning segments of the read were used to extract the corresponding sequence from the original fastq files. The 21-mers in these sequences were counted with Jellyfish [33] and sorted by the counts for each 21-mer in every sample, respectively.

To generate summary statistics for Fig.2, the Breakinator was run on every sample using the “-i in.paf --paf --tabular -o out.txt” parameters. The PacBio scAAV sample was evaluated with the Breakinator using a stricter parameter of 1/4 the default margin (-m 0.025). To determine which breakpoints were recurrent, we extracted the subset of breakpoints that were flagged as either “Foldback”, “Chimeric”, or “True Break”, respectively, then for each file we merged any breakpoints that were within 100 bps of each other to account for inexact alignments around the breakpoint, to determine the number of reads supporting a breakpoint using our merge_breaks.py script. The merging process was run with the -s parameter set to 2, 5, and 10, which sets the minimum number of reads needed to support a breakpoint. The reads that had support of other reads at the respective support thresholds were summed and divided by the total number of reads in the subset to determine the proportion of reads with support, as shown in Fig. 2d.

The SGNEx K562 direct-cDNA replicate 2 run 1 sample was chosen to investigate the raw electrical signal that the ONT base caller interprets because its fast5 files were publicly available, and we discovered numerous putative artifacts in the reads. The fast5 files were indexed with nanopolish using “nanopolish index -d fast5_directory reads.fastq” [34]. Nanopolish eventalign was used to determine which region of the raw signal corresponds to the breakpoint location in the read. To explore the signal in read coordinates, we converted the read fastq file to a fasta file to use as the genome input, then aligned the reads to themselves with “minimap2 -ax map-ont -o reads2reads.sam reads.fastq reads.fa” and used this as the BAM input for the eventalign command as follows: “nanopolish eventalign -r reads.fastq -b reads2reads.bam -g reads.fasta --signal-index”. We chose eight reads in total to investigate the raw signal. We investigated foldbacks, chimeras, and true breakpoints with and without unaligned sequences in the middle, as well as a read with no breakpoint. The start and end of the breakpoint locations in the read were determined from the Breakinator output and converted to the signal coordinates using the nanopolish eventalign output. The fast5 samples were graphed, and the breakpoint coordinates in the signal were highlighted to manually investigate the signal for any signs of increased noise around the breakpoint.

## Results

The Breakinator tool identifies putative foldback and chimeric artifacts by parsing a read alignment file. Our tool extracts the primary and all supplemental alignments of a read and classifies them as either a true breakpoint, chimeric, or foldback using the classification scheme in Fig. 1a. While all metrics can be set by the user, we classify a chimeric artifact as a read that has segments mapping to different chromosomes, or more than 1 Mb apart within the same chromosome. We classify a read as a foldback artifact if the supplementary alignment maps to the same location as the primary alignment (within 200 bps) on the opposite strand. To account for samples where true foldback events may be expected, such as in cancers, we added a symmetry filter to our tool. This filter reports reads as foldback artifacts only if the foldback occurs within a 10% margin on either side of the read’s midpoint, thereby retaining true foldback event-supporting reads.

**Fig. 1.**
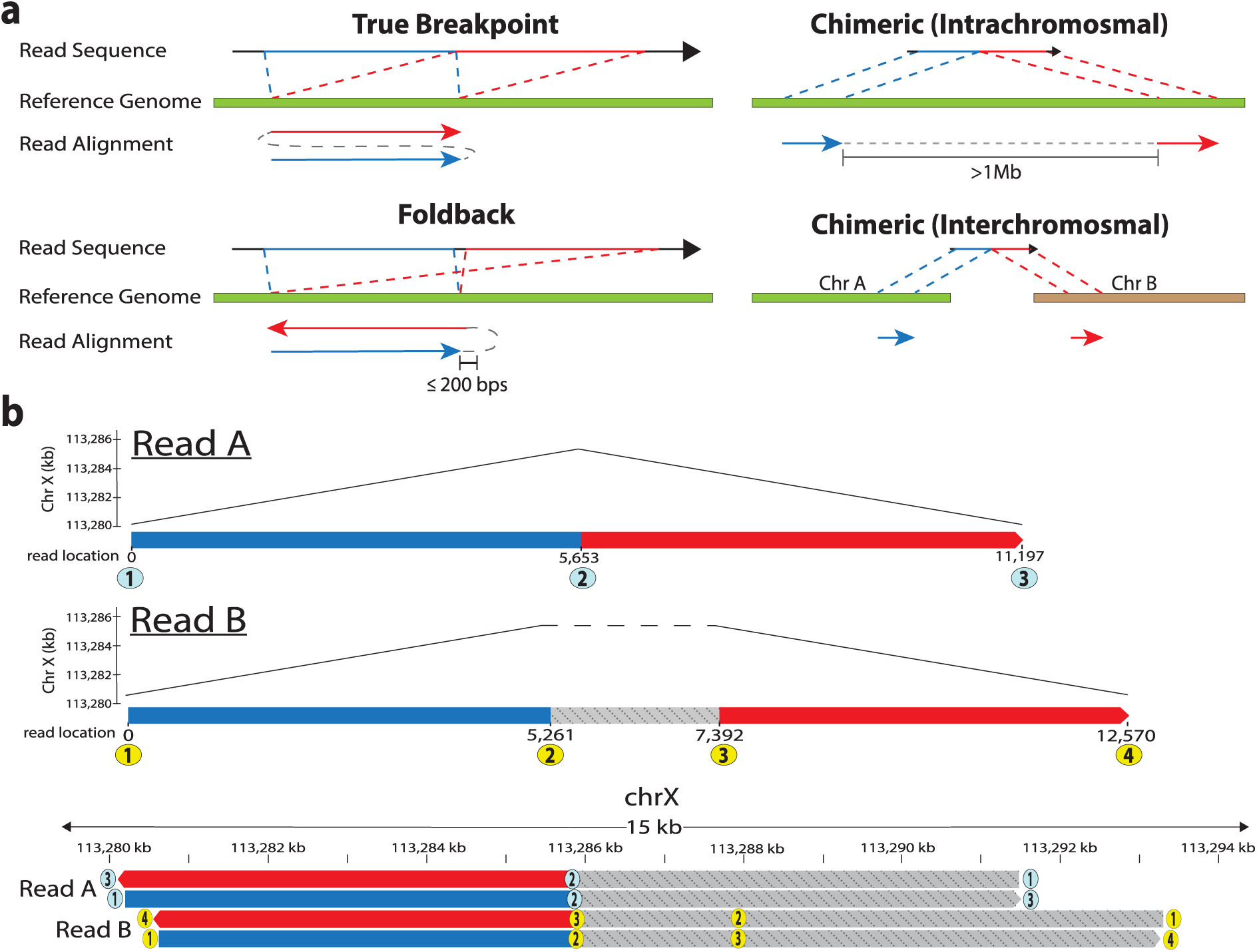
a). Metrics used to define types of breakpoints observed in this study. A “True Break” is a breakpoint that is not found to be chimeric or a foldback, such as the tandem duplication example shown here. A “Foldback” is a read where the second segment maps to approximately the same location (within 200bps) as the first segment, but on the opposite strand. A “Chimeric” is where the second segment maps more than 1 Mb away on the same chromosome (intrachromosomal) or to a different chromosome (interchromosomal). b) IGV schematization of two foldback artifact reads from the HG002_PAW70337_sup sample supporting the same breakpoint at approximately 113,286 kb on chromosome X of the HG002 diploid assembly. Read A (5f80430c-2b83-408b-89b2-63b61b1173cc) is an example of a read with no sequence between the two segments of the foldback. Read B (115c006d-16a6-47b1-bf5c-c5eb9bb85f69) is an example where there are ∼2kb of sequence between the segments.

To verify that our tool properly identifies foldback events, we used PacBio’s samples of self-complementary adeno-associated viruses (scAAV) CBA-eGFP as a positive control. Since scAAV-CBA-eGFP constructs were designed to be perfectly palindromic, we expected our tool to detect them as foldbacks. With the Breakinator, we find that 97.49% CBA-eGFP sample reads are scAAV compared to the 97.69% reported by Talevich *et al*. [23]. This verification also presented an interesting additional use case for the Breakinator for the rapid identification of scAAVs in a sample.

We first noticed a potential technical artifact when analyzing transcriptomic data of the HCC1395 cell line generated by Cotto *et al.* [18]. In the ONT direct-cDNA samples, we observed many reads with supplementary alignments overlapping the primary alignment on the opposite strand, indicative of a foldback SV, that were not present in the ONT direct-RNA sample run on the same RNA replicate. Manual inspection of the unique breakpoint locations in IGV [35] revealed that those detected from direct-RNA had supporting evidence from PacBio HiFi gDNA sequences, whereas most of the direct-cDNA breakpoints did not. This led us to suspect that many of these events were caused by a technical artifact. After developing Breakinator, we used it to identify all breakpoints in the sample and merge breakpoints in the same region (see Methods). We found 35,843 unique breakpoint locations with at least five supporting reads in the direct-cDNA samples, compared to just eight in the direct-RNA. While foldback SVs occur in cancer genomes, they have not been observed at such a high frequency [12], and are not expected to vary so greatly by library type.

To explore whether this was a dataset-specific sequencing result or a systemic technical artifact, we downloaded publicly available datasets representing four library types: direct-RNA, direct-cDNA, cDNA, and gDNA, from four sample types (HG002 cell line, K562 cell line, HCC1395 cell line, and mouse tissue). A range of sequencing chemistries, machines, and base-calling software were evaluated through these samples (Additional file 1). The HG002 samples offer a useful neutral background to investigate technical artifacts because they have a high-quality personal diploid reference assembly to which the reads can be aligned [27]. Therefore, we would expect no germline variants, given that this is the reference, only variations between samples of the cell line, which are likely limited [36]. Each sample was aligned to the corresponding reference genome with the appropriate Minimap2 [3] parameters for each data type (see Methods). Fig. 1b demonstrates two-foldback reads identified by the Breakinator. The reads are both nearly perfectly palindromic and map to the same breakpoint location, yet Read A has no sequence between mappings, while Read B has ∼2kb of sequence that could be searched for retained adaptors. 27 other reads (not shown) cover this foldback location, with no indication of a breakpoint.

Commonly used quality control tools for long reads, such as fastplong, Porechop, pychopper, and Restrander [29–32], did not identify many of the foldback or chimeric reads flagged by the Breakinator (Additional file 2). We found that typically fewer than half of the putative artifactual reads we detected had more than 11 bps of non-mapping sequence between alignments (Additional file 3), making it unlikely that existing tools could detect retained adapter sequences in these reads. An analysis of all 21-mers in the non-mapping sequence between alignments in the k562 samples found these regions enriched in homopolymer, dinucleotide, or trinucleotide repeats. Homopolymer A and the dinucleotide AC repeat were observed as the most frequent sequences in our analysis.

Summary statistics across library types and samples evaluated with the Breakinator are shown in Fig. 2a. ONT direct-cDNA samples were found to have substantially elevated levels of foldback artifacts (∼10-20% of all reads). Meanwhile, at most one foldback read was typically observed in each of the ONT direct-RNA samples. The ONT cDNA samples had the lowest rates of foldback artifacts (∼0.05 – 0.10%). In ONT gDNA, we observed consistent foldback rates of ∼0.55–0.6%. While only three PacBio samples were investigated, they did not show any evidence of systemic foldback artifacts. There were virtually no foldback reads observed in the PacBio HiFi (Revio) or Mas-Seq (Vega) data, while 0.03% of the Iso-Seq (Sequel II) reads were considered foldbacks. For five of the samples, from K562 and HCC1395 cells, the same RNA replicate was sequenced with different technologies, allowing for direct comparisons in the rates of foldbacks and chimeras across libraries without replicate bias. It was observed that for all replicates, the rates of foldbacks and chimeras were much higher in ONT direct-cDNA library samples than in other libraries, as shown in Fig. 2b.

**Fig. 2.**
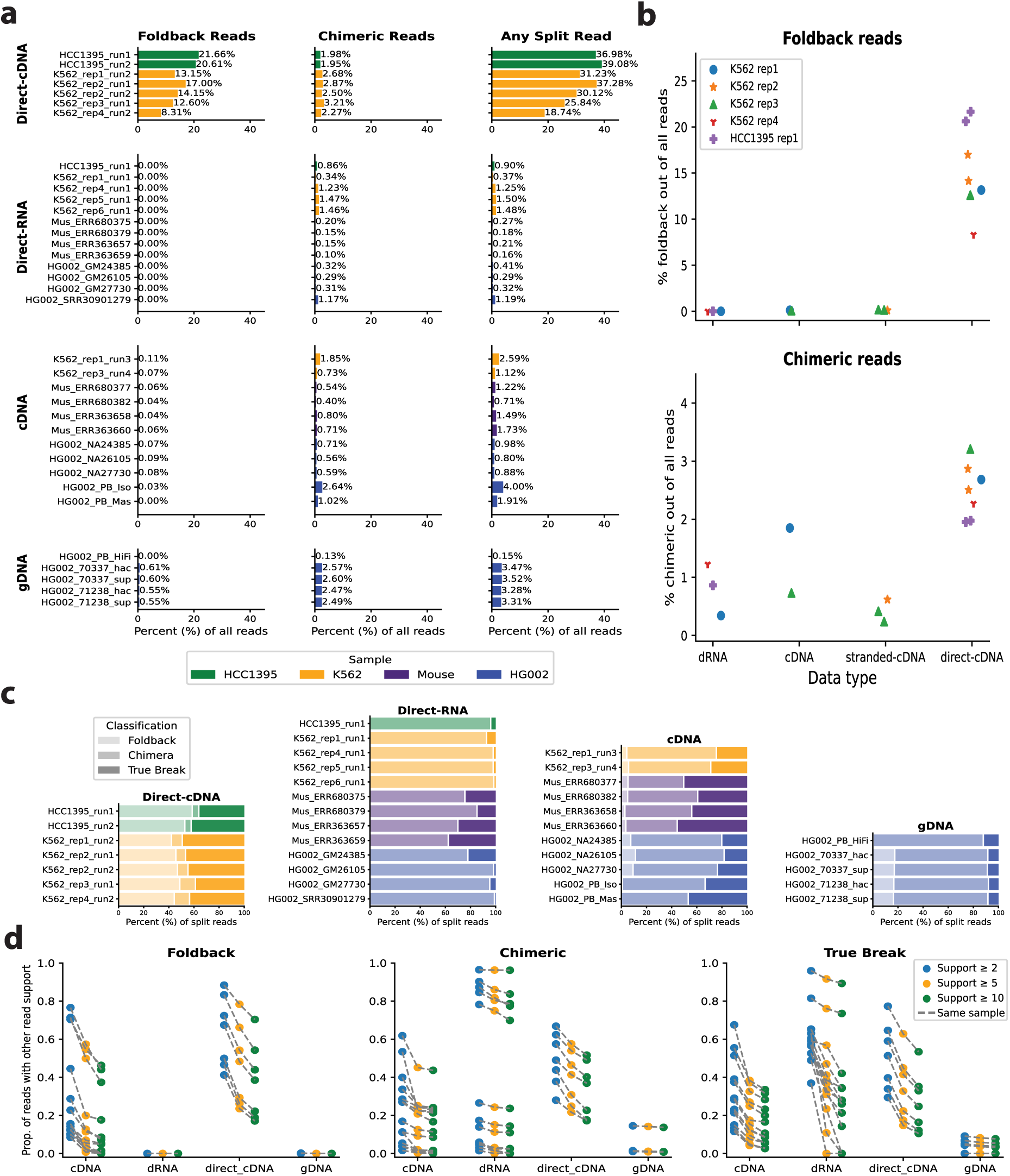
a) The observed rates of Foldback, Chimeric, and any split read out of all reads that passed the filtering criteria for direct-cDNA, direct-RNA, cDNA, and gDNA data from four different sample types: HCC1395, K562, mouse tissue, and HG002. B) Paired comparisons for the five RNA replicates that were sequenced with different technologies for both foldback and chimeric reads. c) The proportion of Foldback, Chimeric, and True Break reads in relation to the number of split reads. d) Recurrence of breakpoints at 2, 5, and 10 supporting reads for Foldback, Chimeric, and True Break reads.

To explore the potential effects of these artifactual reads in SV calling, we investigated the proportion of reads used for SV calling, namely, reads with split alignments, that they represent. In cDNA, foldback artifacts account for 5-15% of all split reads, while chimeras are ∼30-50%. In gDNA, foldbacks are also ∼18% of split reads, and chimeric reads are ∼70% of all split reads (Fig. 2.c). These artifactual reads were also observed to be recurrent at the same locations, which made it more likely that they would be called as false positive SVs. Fig. 2d highlights that many of the locations to which these reads align have more than 2, 5, or 10 reads of support. More recent transcriptomic data from the HG002 cell line comprise the cluster of higher proportion samples in the cDNA foldback and dRNA chimeric evaluations in Fig. 2d. We also observed that the ONT HG002 gDNA data, which was generated on the latest R10.4 pores, had few recurrent foldback locations, typically with only two supporting reads. Finally, we investigated the raw signal of eight putative artifactual reads from the SG-NEx K562 direct-cDNA replicate 2 run 1 to visualize the electrical signal interpreted by the base-caller. We did not observe any notable difference in signal around the breakpoint region for artifacts compared to true breakpoints (Additional file 4).

## Discussion

In this study, we documented the rate of foldback artifacts in long-read sequencing data across a diverse set of samples. In addition, we developed the open-source Breakinator tool to report putative foldback artifacts, as well as the already well-characterized chimeric artifacts, in long-read data. While an alignment-based artifact detection method, as is implemented by the Breakinator, may lead to issues in distinguishing true SV reads from artifact reads and introduce reference bias, the increased specificity offered by alignment to a reference can be useful; This is evidenced by our ability to detect foldback artifact reads that are missed by commonly used quality control tools such as fastplong, Porechop, Restrander, and pychopper [29–32]. We suspect these artifacts are missed due to the tools’ reliance on identifying adapter sequences within the middle of a read sequence to identify a concatemerized read. We find that less than half of the putative artifactual reads identified by the Breakinator have any searchable sequence between alignments of the read segments (Fig. 1b). Various assemblers, such as Canu, Flye, and MECAT, use the alignments of reads to each other or assembled contigs to detect technical artifacts [37–39]. This approach is free of reference bias and does not rely on adaptor sequence matches, yet it is computationally expensive and does not scale well to a quality control tool that would not otherwise perform these alignments.

While the rates of putative foldback and chimeric artifacts across all reads are relatively low for libraries other than ONT direct-cDNA (Fig. 2a), they represent a far larger proportion of the split alignment reads used for SV calling. Fig. 2c highlights that these artifacts can cause issues even with ONT gDNA or cDNA data, in which foldback artifacts represent at most 0.61% of all reads, yet represent approximately 15% of the split reads used for SV calling. We suspect that the rate of foldback artifacts is lowest in the cDNA samples, as cDNA is the only library type in this study that undergoes PCR amplification. Palindromic sequences, such as foldbacks, are known to be difficult to PCR amplify, as they can bind to themselves [40, 41].

Given that the rates of foldback artifacts appear relatively consistent among sequencing runs of the same data type, but vary between data types that are prepared with different chemistries, we believe it is most likely that these artifacts arise during library preparation, rather than a failure of the base-caller to recognize a pore-open signal or an adapter sequence, as was hypothesized by Soneson *et al*. [11]. Additionally, we observed both forward and reverse strands of foldback and chimeric reads, which can only arise during library preparation, compared to an informatic chimera, which would only occur in one reading direction. Lastly, our analysis of a subset of ONT raw electrical signals did not reveal any noise around putative breakpoints, indicating that the foldback artifact molecules likely passed through the pore without issue.

We envision that Breakinator can serve as either a filtering or diagnostic tool, depending on the sample and data type. In normal samples, both transcriptomic and genomic, we expect most flagged reads to represent true artifacts and would recommend they be filtered before downstream analysis of the alignment files. Flagged foldback reads can be handled in multiple ways, such as removing them completely, retaining only their primary alignment, or splitting them into unique alignments. A limitation of our approach is the difficulty in distinguishing real somatic events from artifacts in gDNA cancer samples, where large-scale rearrangements and foldback events are common [12]. In these cases, a more nuanced diagnostic approach may be appropriate, such as inspecting structural variant calls based on the Breakinator’s classification of their supporting reads. However, if a read is nearly perfectly palindromic, with a breakpoint occurring near the center as identified by our symmetry filter, we believe it can reasonably be filtered out. If the event is real, there should still be sufficient supporting reads that are not perfectly palindromic for the event.

## Conclusions

We present evidence of an unresolved technical artifact, a foldback read. If these technical artifacts are not properly removed, they can lead to tangles in an assembly graph or false-positive SV calls. We developed the open-source Breakinator tool to flag putative foldback and chimeric technical artifacts in long-read sequences. Many current quality control tools rely on detecting incorrectly incorporated adapter sequences, but through a reference alignment-based approach, we can detect reads that may not have retained non-sample sequence, but still likely represent a non-real biological concatenation. In this study, we profiled both ONT and PacBio data across a range of specimens, library types, sequencing chemistries, sequencing machines, and base-callers. We find that foldback artifacts occur throughout ONT library types and are particularly high in direct-cDNA libraries. We believe the Breakinator will offer a useful tool for quality control in long-read sequencing analyses, particularly for SV calling.

## Declarations

### Ethics approval and consent to participate

Not applicable.

### Consent for publication

Not applicable.

### Additional files

Additional_file_1.xlsx – All public data used in this study and their sources. Includes metadata on datatype, sequencer, pore, chemistry, and base-caller.

Additional_file_2.xlsx – Proportions of reads remaining, including artifact subsets, after filtering with other current tools (fastplong, Porechop, pychopper, and Restrander).

Additional_file_3.xlsx – Proportions of reads that had 11 bps or greater of unmapped read sequence between alignment blocks.

Additional_file_4.html – SGNEx K562 direct-cDNA replicate 2 run 1 raw signal analysis around identified breakpoints for eight reads.

### Availability of data and materials

The Breakinator tool and additional software are available on GitHub at https://github.com/jheinz27/breakinator. All data used in this study were previously made publicly available, and their sources are listed in Additional file 1. The alignment files for every sample, as well as all breakpoints identified and their Breakinator classification, are available on Zenodo at: https://zenodo.org/records/17567592.

### Competing interests

M.M. holds equity in Bayer, Delve Bio, Isabl, and Karyoverse; consults for Delve Bio; receives research funding from Bayer; and receives patent licensing payments from Bayer and Labcorp.

### Funding

J.M.H. is supported by the U.S. National Institutes of Health training grant T32HG002295. M.M. is supported by NIH grants R35CA197568, U24CA264029, and U24CA294203. H.L. is supported by NIH grants U24CA294203 and R01HG010040.

### Authors’ contributions

J.M.H., M.M., and H.L conceived the project. J.M.H. implemented the method, analyzed the data, and drafted the manuscript. M.M. and H.L provided edits. All authors read and approved the final manuscript.

## Acknowledgments

We would like to thank Drs. Neng Huang, Owen Hirschi, Alaina Shumate, Maximillian Marin, Victoria Popic, and Kar-Tong Tan, as well as all members of the Meyerson and Li labs, for their helpful discussions.

## Notes

https://github.com/jheinz27/breakinator

https://zenodo.org/records/17567592

### Summary of Updates

Updated our method, Breakinator, to support the more common SAM/BAM/CRAM format files along with PAF files, highlighted that sequence between aligning read segments in artifact reads is enriched for repetitive elements, and added a paragraph discussing the envisioned use case of Breakinator in an analysis pipeline depending on sample and datatype.

