## Additional File 4 for "Detecting Foldback Artifacts in Long-reads": Additional_file_4.html

raw\_signal


In [2]:

```
import h5py
import matplotlib.pyplot as plt
import numpy as np
```

The raw signal of 8 reads from SGNEx K562 direct-cDNA replicate 2 run 1 (link) were investigated in greater detail. Foldback, Chimeric, True Break, and no Break reads are shown before. The region that corresponds to the breakpoint location in the read as identified by the Breakinator is denoted by the dashed red lines below. The entire signal of the read is shown as well as zoomed in window for around the breakpoint for every split read.

### 0010753f-4cf6-46b4-a6d6-2f355a84701a (foldback no seq in middle)¶

In [ ]:

```
0010753f-4cf6-46b4-a6d6-2f355a84701a    1158    221     653     -       NC_060926.1     242696752       130808877       130809308       423     438     60      tp:A:P  mm:i:2  gn:i:13 go:i:7  cg:Z:11M2D10M2D4M4I57M1I14M1D160M2I93M1D76M

0010753f-4cf6-46b4-a6d6-2f355a84701a    1158    645     1000    +       NC_060926.1     242696752       130808942       130809308       351     369     60      tp:A:P  mm:i:1  gn:i:17 go:i:10 cg:Z:68M1D4M2D1M1D108M1D29M1I5M1I15M1I4M1D10M1D69M7D39M
```

In [9]:

```
fast5_path = 'GXB01130_20181017_FAK11045_GA30000_sequencing_run_171018_K562_mRNA_directcDNA_88106_read_878_ch_224_strand.fast5'

with h5py.File(fast5_path, 'r') as f:
    reads  = f['Raw/Reads']
    read_id, = reads.keys()          
    signal = reads[read_id]['Signal'][:] 

plt.figure(figsize=(12,4))
plt.plot(signal, linewidth=1)
plt.axvline(x=7774, color='red', linestyle='--', linewidth=1)
plt.axvline(x=7885, color='red', linestyle='--', linewidth=1) 
plt.title(f'Raw ONT signal for read 0010753f-4cf6-46b4-a6d6-2f355a84701a')
plt.xlabel('Sample index')
plt.ylabel('Digital signal amplitude')
plt.tight_layout()
plt.show()

with h5py.File(fast5_path, 'r') as f:
    reads  = f['Raw/Reads']
    read_id, = reads.keys()           
    signal = reads[read_id]['Signal'][7000:8500]  

plt.figure(figsize=(12,4))
plt.plot(list(range(7000,8500)),signal, linewidth=1)
with open('002c731f-019b-495c-bb55-e680b63e7194_eventalign.tsv','r') as f: 
    for line in f:
        s = int(line.strip().split('\t')[-1])
        if 7000< s < 8500:
            plt.axvline(x=s, color='black', linestyle='--', linewidth=0.1)
plt.axvline(x=7774, color='red', linestyle='--', linewidth=1)
plt.axvline(x=7885, color='red', linestyle='--', linewidth=1)
plt.title(f'Zoomed Raw ONT signal for read 0010753f-4cf6-46b4-a6d6-2f355a84701a')
plt.xlabel('Sample index')
plt.ylabel('Digital signal amplitude')
plt.tight_layout()
plt.show()
```

### 01005106-eb3f-4c9f-bbf0-261e0636177a (foldback seq in middle)¶

(has sequence in middle: AAAAAAAAAAAAAAAAAAAAATAGAGCGAC )

In [ ]:

```
01005106-eb3f-4c9f-bbf0-261e0636177a    1188    60      607     -       NC_060929.1     182045439       150980767       150986271       537     551     60      tp:A:P  mm:i:4  gn:i:10 go:i:8  cg:Z:74M1D44M2I1M1255N12M1I65M1116N162M621N39M1D58M1I25M1D11M1D15M1967N27M2I8M
01005106-eb3f-4c9f-bbf0-261e0636177a    1188    637     1142    +       NC_060929.1     182045439       150980780       150986267       503     528     60      tp:A:P  mm:i:2  gn:i:23 go:i:9  cg:Z:61M1D7M2D36M1255N9M7D7M1D26M1D11M1D13M1D1116N35M6D48M3D70M621N151M1967N31M\
#breakpoint called
NC_060929.1     150980767       <>      NC_060929.1     150980780       60      01005106-eb3f-4c9f-bbf0-261e0636177a    Foldback
```

In [13]:

```
fast5_path = 'GXB01130_20181017_FAK11045_GA30000_sequencing_run_171018_K562_mRNA_directcDNA_88106_read_90181_ch_57_strand.fast5'

with h5py.File(fast5_path, 'r') as f:
    reads  = f['Raw/Reads']
    read_id, = reads.keys()               
    signal = reads[read_id]['Signal'][:] 

plt.figure(figsize=(12,4))
plt.plot(signal, linewidth=1)
plt.axvline(x=8067, color='red', linestyle='--', linewidth=1)
plt.axvline(x=8581, color='red', linestyle='--', linewidth=1)
plt.title(f'Raw ONT signal for read 01005106-eb3f-4c9f-bbf0-261e0636177a')
plt.xlabel('Sample index')
plt.ylabel('Digital signal amplitude')
plt.tight_layout()
plt.show()

with h5py.File(fast5_path, 'r') as f:
    reads  = f['Raw/Reads']
    read_id, = reads.keys()             
    signal = reads[read_id]['Signal'][7500:9000]

plt.figure(figsize=(12,4))
plt.plot(list(range(7500,9000)),signal, linewidth=1)
with open('01005106-eb3f-4c9f-bbf0-261e0636177a_eventalign.tsv','r') as f: 
    for line in f:
        s = int(line.strip().split('\t')[-1])
        if 7500< s < 9000:
            plt.axvline(x=s, color='black', linestyle='--', linewidth=0.1)
plt.axvline(x=8067, color='red', linestyle='--', linewidth=1)
plt.axvline(x=8581, color='red', linestyle='--', linewidth=1)
plt.title(f'Zoomed Raw ONT signal for read 01005106-eb3f-4c9f-bbf0-261e0636177a')
plt.xlabel('Sample index')
plt.ylabel('Digital signal amplitude')
plt.tight_layout()
plt.show()

with h5py.File(fast5_path, 'r') as f:
    reads  = f['Raw/Reads']
    read_id, = reads.keys()            
    signal1 = reads[read_id]['Signal'][:8324] 
    signal2 = reads[read_id]['Signal'][8324:]
```

### 01cf6ed6-8abc-4435-af52-d993bbcd8ca6 (foldback no seq in middle)¶

In [ ]:

```
01cf6ed6-8abc-4435-af52-d993bbcd8ca6    6901    24      3954    +       NC_060933.1     150617247       133055393       133059349       3857    3989    60      tp:A:P  mm:i:40 gn:i:92 go:i:68 cg:Z:7M2I7M1D2M1D20M2D13M3D309M2I183M1I116M1D70M1D8M1I12M1I38M1D28M1D11M3D3M1D49M1I15M1I54M1I52M1D7M2D28M1I24M1D42M1D42M2D68M1I23M4D36M2I57M1D105M1I2M2D35M1I38M1D130M3D63M1D236M1D42M2D89M1D15M1D1M1D62M1D174M1D57M1I10M5D36M1D13M1D15M2D209M1D116M1I13M1I21M1D8M1I7M2D32M1I60M1D63M1D97M1I183M1I50M1D3M1D106M1I61M1I9M1I75M1I77M1I1M2I5M1I89M1I76M1I89M
01cf6ed6-8abc-4435-af52-d993bbcd8ca6    6901    3936    6888    -       NC_060933.1     150617247       133055382       133059349       2516    3774    60      tp:A:P  mm:i:293        gn:i:965        go:i:335        cg:Z:24M2D11M15D13M1D35M1D1M1D22M4I71M8D14M1D6M12D16M9D4M2D3M1D5M1D2M2D3M2D39M5D6M1D4M1D4M106N11M2I6M2I10M1I3M1I3M1D3M1I2M1D5M7I13M1D3M1I9M14D10M1D10M1I26M1I9M1D20M1D17M9D10M7D3M2D2M6D5M1I6M1D8M6D17M3D6M2D2M3D2M1I2M1I9M1I14M2I2M1I19M1I4M1D9M1D5M11D1M1D4M10D4M2D1M1D3M1D2M3D3M2D25M3D16M3I6M1I22M1I1M1I4M2I13M2I3M1I13M1D30M3D5M6D16M4D7M3D1M1D4M2D1M1D21M1D28M2I12M1I5M2I10M2I9M4D5M12D2M4D5M1I13M1I15M1D6M1D14M2I2M1D2M2I10M1D9M4D2M2I10M1I18M1D15M6D14M2D3M2D3M2D4M1I3M1D2M2I6M1D3M1D50M7D19M14D12M1D8M2D1M5D4M2I3M1D11M1D5M1I3M3D3M3D22M6D3M1D11M2D3M1D9M1D11M1D1M13D6M1D7M3D4M8D12M1D24M1D3M3D3M3D2M3D2M1D8M2D6M10D1M1D5M1D2M4D2M5D1M4D6M10D2M5D13M1D17M2D17M3D16M5D18M4I4M3D6M1D19M1D8M14D9M1D7M17D4M2D10M1D5M1D16M3D12M1D1M1D5M1D1M4D6M6D2M1D6M1D7M7D5M5D7M3D7M1I34M2I3M7D8M1D2M1I19M1D28M1D8M6D4M2D21M1D23M1I4M2D5M4D16M1D1M6D18M3D6M2D1M1D1M1D5M6D3M2D5M13D2M3D10M2D5M1D15M3D2M2D1M1D6M4D2M4D2M3D2M11D1M1D16M1D9M3I8M1I5M2D6M1D2M3D14M1D8M3D3M1D4M1D6M2I3M1D35M1D4M1D1M3D20M10D2M2D8M1D8M7D11M1I5M5D20M3D3M3I4M230N10M1I2M2D3M2D2M1D10M1I8M2I2M1I8M2D8M2I4M1I6M9I5M1D1M1D8M1I16M1I5M2I5M2I7M4I1M2D8M1D5M1I4M1I2M1I4M5I5M5D1M1D7M6D15M5D7M4D4M1D4M1I3M2D8M5D4M1D2M1D5M1D5M1D3M8D17M8D5M4D2M4D8M4D7M1D18M3D10M1I10M1I6M2I21M1D3M3D5M2D1M1D4M4D1M3D5M1I11M1I10M1I26M2D3M3D6M8D1M1D10M1I10M2I9M1I5M1D6M1D7M4D5M6D3M1D10M2D4M19D1M1D4M4D9M11D14M1I1M1I7M2D12M2I12M1D19M1D12M6D13M1D1M1D2M4D7M6D1M1D2M4D5M4D9M1I2M1D3M2D5M5D19M2I8M3D10M4D10M1D4M1D7M2D1M1D6M1D9M3D16M3I15M1D15M3I26M6D15M1D3M1D6M3D6M13D11M2D14M3D17M1I8M1I5M1D6M
NC_060933.1     133059349       ><      NC_060933.1     133059349       60      01cf6ed6-8abc-4435-af52-d993bbcd8ca6    Foldback
```

In [15]:

```
fast5_path = 'GXB01130_20181017_FAK11045_GA30000_sequencing_run_171018_K562_mRNA_directcDNA_88106_read_56908_ch_140_strand.fast5'

with h5py.File(fast5_path, 'r') as f:
    reads  = f['Raw/Reads']
    read_id, = reads.keys()             
    signal = reads[read_id]['Signal'][:] 

plt.figure(figsize=(12,4))
plt.plot(signal, linewidth=0.3)
plt.axvline(x=45848, color='red', linestyle='--', linewidth=1)
plt.axvline(x=46045, color='red', linestyle='--', linewidth=1)
plt.title(f'Raw ONT signal for 01cf6ed6-8abc-4435-af52-d993bbcd8ca6')
plt.xlabel('Sample index')
plt.ylabel('Digital signal amplitude')
plt.tight_layout()
plt.show()

r1 = 45000
r2 = 47000
with h5py.File(fast5_path, 'r') as f:
    reads  = f['Raw/Reads']
    read_id, = reads.keys()               # unpack the single read ID
    signal = reads[read_id]['Signal'][r1:r2]  # raw digital trace

plt.figure(figsize=(12,4))
plt.plot(list(range(r1,r2)),signal, linewidth=1)
with open('01cf6ed6-8abc-4435-af52-d993bbcd8ca6_eventalign.tsv','r') as f: 
    for line in f:
        s = int(line.strip().split('\t')[-1])
        if r1< s < r2:
            plt.axvline(x=s, color='black', linestyle='--', linewidth=0.1)
plt.axvline(x=45848, color='red', linestyle='--', linewidth=1)
plt.axvline(x=46045, color='red', linestyle='--', linewidth=1)
plt.title(f'Zoomed Raw ONT signal for read 01cf6ed6-8abc-4435-af52-d993bbcd8ca6')
plt.xlabel('Sample index')
plt.ylabel('Digital signal amplitude')
plt.tight_layout()
plt.show()
```

### 00eb9e4b-271f-4245-ad7b-0a4c84762077 (chimeric no seq in middle)¶

In [ ]:

```
00eb9e4b-271f-4245-ad7b-0a4c84762077    1330    256     986     -       NC_060939.1     99753195        28736432        28737186        710     761     60      tp:A:P  mm:i:13 gn:i:38 go:i:20 cg:Z:38M1I32M1I21M3I32M1D20M1I59M1D55M3D22M1D4M1D63M1D8M1I34M1D20M1D21M1D1M1D15M1D151M12D72M1D17M4D4M1D34M
00eb9e4b-271f-4245-ad7b-0a4c84762077    1330    976     1261    +       NC_060945.1     45090682        37576897        37577199        277     305     60      tp:A:P  mm:i:5  gn:i:23 go:i:13 cg:Z:36M3D2M3D29M1D28M3D16M1I20M2D6M1D42M1I60M1D25M1I3M3D1M1D1M2D13M
NC_060939.1     28736432        <>      NC_060945.1     37576897        60      00eb9e4b-271f-4245-ad7b-0a4c84762077    Chimeric
```

In [17]:

```
fast5_path = 'GXB01130_20181017_FAK11045_GA30000_sequencing_run_171018_K562_mRNA_directcDNA_88106_read_32414_ch_294_strand.fast5'

with h5py.File(fast5_path, 'r') as f:
    reads  = f['Raw/Reads']
    read_id, = reads.keys()            
    signal = reads[read_id]['Signal'][:]  

plt.figure(figsize=(12,4))
plt.plot(signal, linewidth=1)
plt.axvline(x=8637, color='red', linestyle='--', linewidth=1)
plt.axvline(x=8717, color='red', linestyle='--', linewidth=1)
plt.title(f'Raw ONT signal for read 00eb9e4b-271f-4245-ad7b-0a4c84762077')
plt.xlabel('Sample index')
plt.ylabel('Digital signal amplitude')
plt.tight_layout()
plt.show()

with h5py.File(fast5_path, 'r') as f:
    reads  = f['Raw/Reads']
    read_id, = reads.keys()               
    signal = reads[read_id]['Signal'][7500:9000]  

r1 = 8000
r2 = 9000
with h5py.File(fast5_path, 'r') as f:
    reads  = f['Raw/Reads']
    read_id, = reads.keys()              
    signal = reads[read_id]['Signal'][r1:r2]

plt.figure(figsize=(12,4))
plt.plot(list(range(r1,r2)),signal, linewidth=1)
with open('00eb9e4b-271f-4245-ad7b-0a4c84762077_eventalign.tsv','r') as f: 
    for line in f:
        s = int(line.strip().split('\t')[-1])
        if r1< s < r2:
            plt.axvline(x=s, color='black', linestyle='--', linewidth=0.1)
plt.axvline(x=8637, color='red', linestyle='--', linewidth=1)
plt.axvline(x=8717, color='red', linestyle='--', linewidth=1)
plt.title(f'Zoomed Raw ONT signal for read 00eb9e4b-271f-4245-ad7b-0a4c84762077')
plt.xlabel('Sample index')
plt.ylabel('Digital signal amplitude')
plt.tight_layout()
plt.show()
```

### 04532c71-be2f-452f-814d-0fbab91f01f0 (chimeric seq in middle)¶

In [ ]:

```
04532c71-be2f-452f-814d-0fbab91f01f0    2785    111     1602    +       NC_060936.1     133324548       52861777        52869379        1451    1518    60      tp:A:P  mm:i:28 gn:i:39 go:i:24 cg:Z:33M2D15M1D106M2I1M1D32M1D32M3I145M6D53M841N56M1I3M159N5M1I2M1I201M1D12M875N132M1I52M1D36M1D32M1D1M1D34M522N15M1D7M4D2M3D3M1I9M2D50M475N61M644N123M1I24M1I16M1D45M2580N141M
04532c71-be2f-452f-814d-0fbab91f01f0    2785    1624    2742    +       NC_060932.1     146259331       62002190        62003676        947     1400    60      tp:A:P  mm:i:131        gn:i:322        go:i:115        cg:Z:17M6D10M28D23M2I18M1I14M1D11M1I10M1I9M126N6M1D10M1D8M1D6M1I18M2D11M2I23M6D10M2D2M1D5M6D3M1D7M1D13M1D21M2D4M1D7M6D2M1D3M1D6M1D8M2D3M1D10M1D19M3D9M2D2M2D16M2I10M1D3M2D10M2D6M7D8M2D13M2I12M3I13M3I2M1D21M1D15M5D1M3D1M1D14M11D3M2D2M8D1M3D2M1D5M7D10M1D15M1I17M1D13M6D12M1D2M2D11M1D3M1D3M2D5M6D14M3I7M1D13M1I4M1D1M1D12M13D2M1D13M1D26M5D34M1D16M1D2M2I17M1D3M7D3M3D11M1I4M1D2M5D10M1D7M2D2M1I2M1D5M2D8M1D10M3D4M3D2M9D1M1D13M2D11M1I8M2D7M2D3M6I1M1D6M1D2M1D2M2D2M1D2M1D10M1I6M1I11M2D3M5D15M1D11M1I12M1I3M1D19M8D26M5D11M7D15M12D3M2I40M4D24M
NC_060936.1     52869379        >>      NC_060932.1     62002190        60      04532c71-be2f-452f-814d-0fbab91f01f0    Chimeric
```

In [19]:

```
fast5_path = 'GXB01130_20181017_FAK11045_GA30000_sequencing_run_171018_K562_mRNA_directcDNA_88106_read_29384_ch_420_strand.fast5'

with h5py.File(fast5_path, 'r') as f:
    reads  = f['Raw/Reads']
    read_id, = reads.keys()               
    signal = reads[read_id]['Signal'][:] 

plt.figure(figsize=(12,4))
plt.plot(signal, linewidth=0.5)
plt.axvline(x=16836, color='red', linestyle='--', linewidth=1)
plt.axvline(x=17020, color='red', linestyle='--', linewidth=1)
plt.title(f'Raw ONT signal for read 04532c71-be2f-452f-814d-0fbab91f01f0')
plt.xlabel('Sample index')
plt.ylabel('Digital signal amplitude')
plt.tight_layout()
plt.show()

with h5py.File(fast5_path, 'r') as f:
    reads  = f['Raw/Reads']
    read_id, = reads.keys()              
    signal = reads[read_id]['Signal'][7500:9000]  

r1 = 16000
r2 = 18000
with h5py.File(fast5_path, 'r') as f:
    reads  = f['Raw/Reads']
    read_id, = reads.keys()               
    signal = reads[read_id]['Signal'][r1:r2]  

plt.figure(figsize=(12,4))
plt.plot(list(range(r1,r2)),signal, linewidth=1)
with open('04532c71-be2f-452f-814d-0fbab91f01f0_eventalign.tsv','r') as f: 
    for line in f:
        s = int(line.strip().split('\t')[-1])
        if r1< s < r2:
            plt.axvline(x=s, color='black', linestyle='--', linewidth=0.1)
plt.axvline(x=16836, color='red', linestyle='--', linewidth=1)
plt.axvline(x=17020, color='red', linestyle='--', linewidth=1)
plt.title(f'Zoomed Raw ONT signal for read 04532c71-be2f-452f-814d-0fbab91f01f0')
plt.xlabel('Sample index')
plt.ylabel('Digital signal amplitude')
plt.tight_layout()
plt.show()
```

### 074d4e1a-4709-4b9e-8dc5-407190dd006f (true breakpoint example)¶

In [ ]:

```
074d4e1a-4709-4b9e-8dc5-407190dd006f    2968    1079    1845    -       NC_060927.1     201105948       145767880       145775648       748     773     60      tp:A:P  mm:i:10 gn:i:15 go:i:10 cg:Z:54M656N33M90N24M2D57M941N16M1I77M2D57M2I14M190N56M1I63M1D34M1D33M1D23M910N5M2I39M3130N105M1086N22M2I46M
074d4e1a-4709-4b9e-8dc5-407190dd006f    2968    64      1077    -       NC_060927.1     201105948       145748862       145769904       998     1031    60      tp:A:P  mm:i:8  gn:i:25 go:i:18 cg:Z:53M9140N27M1D17M1362N115M1I17M1957N1M2D15M1I71M1I10M1847N2I21M1I19M1D74M470N18M2D43M1D9M1D60M2993N11M2D9M2D3M1D40M562N33M1I51M2D9M656N33M90N83M941N8M2D142M1D14M
NC_060927.1     145748862       <<      NC_060927.1     145775648       60      074d4e1a-4709-4b9e-8dc5-407190dd006f    Pass
```

In [21]:

```
fast5_path = 'GXB01130_20181017_FAK11045_GA30000_sequencing_run_171018_K562_mRNA_directcDNA_88106_read_18360_ch_146_strand.fast5'

with h5py.File(fast5_path, 'r') as f:
    reads  = f['Raw/Reads']
    read_id, = reads.keys()             
    signal = reads[read_id]['Signal'][:] 

plt.figure(figsize=(12,4))
plt.plot(signal, linewidth=1)
plt.axvline(x=10668, color='red', linestyle='--', linewidth=1)
plt.axvline(x=10696, color='red', linestyle='--', linewidth=1)
plt.title(f'Raw ONT signal for read 074d4e1a-4709-4b9e-8dc5-407190dd006f')
plt.xlabel('Sample index')
plt.ylabel('Digital signal amplitude')
plt.tight_layout()
plt.show()

with h5py.File(fast5_path, 'r') as f:
    reads  = f['Raw/Reads']
    read_id, = reads.keys()          
    signal = reads[read_id]['Signal'][7500:9000]  

r1 = 9500
r2 = 11500
with h5py.File(fast5_path, 'r') as f:
    reads  = f['Raw/Reads']
    read_id, = reads.keys()               
    signal = reads[read_id]['Signal'][r1:r2]  

plt.figure(figsize=(12,4))
plt.plot(list(range(r1,r2)),signal, linewidth=1)
with open('074d4e1a-4709-4b9e-8dc5-407190dd006f_eventalign.tsv','r') as f: 
    for line in f:
        s = int(line.strip().split('\t')[-1])
        if r1< s < r2:
            plt.axvline(x=s, color='black', linestyle='--', linewidth=0.1)
plt.axvline(x=10668, color='red', linestyle='--', linewidth=1)
plt.axvline(x=10696, color='red', linestyle='--', linewidth=1)
plt.title(f'Zoomed Raw ONT signal for read 074d4e1a-4709-4b9e-8dc5-407190dd006f')
plt.xlabel('Sample index')
plt.ylabel('Digital signal amplitude')
plt.tight_layout()
plt.show()
```

### 000f8add-2084-4768-9c05-a27e4321ff9c (no breakpoint passes filter)¶

In [23]:

```
fast5_path = 'GXB01130_20181017_FAK11045_GA30000_sequencing_run_171018_K562_mRNA_directcDNA_88106_read_9075_ch_193_strand2.fast5'

with h5py.File(fast5_path, 'r') as f:
    reads  = f['Raw/Reads']
    read_id, = reads.keys()               
    signal = reads[read_id]['Signal'][:]  

plt.figure(figsize=(12,4))
plt.plot(signal, linewidth=1)
plt.title(f'Raw ONT signal for read 000f8add-2084-4768-9c05-a27e4321ff9c')
plt.xlabel('Sample index')
plt.ylabel('Digital signal amplitude')
plt.tight_layout()
plt.show()
```

### 4daaec37-da49-4578-b710-5cb70e0083e9 (no breakpoint passes filter)¶

In [25]:

```
fast5_path = 'GXB01130_20181017_FAK11045_GA30000_sequencing_run_171018_K562_mRNA_directcDNA_88106_read_6506_ch_259_strand.fast5'

with h5py.File(fast5_path, 'r') as f:
    reads  = f['Raw/Reads']
    read_id, = reads.keys()               
    signal = reads[read_id]['Signal'][:]  

plt.figure(figsize=(12,4))
plt.plot(signal, linewidth=1)
plt.title(f'Raw ONT signal for read 000f8add-2084-4768-9c05-a27e4321ff9c')
plt.xlabel('Sample index')
plt.ylabel('Digital signal amplitude')
plt.tight_layout()
plt.show()
```

In [ ]:
